# Higher-order patterns of aquatic species spread through the global shipping network

**DOI:** 10.1101/704684

**Authors:** Mandana Saebi, Jian Xu, Erin K. Grey, David M. Lodge, Nitesh Chawla

## Abstract

The introduction and establishment of non-indigenous species (NIS) through global ship movements is a significant threat to marine ecosystems and economies. While ballast-vectored invasions have been partly addressed by some national policies and an international agreement regulating the concentrations of organisms in ballast water, biofouling-vectored invasions remain a large risk. Development of additional realistic and cost-effective ship-borne NIS policies requires an accurate estimation of NIS spread risk from both ballast water and biofouling. In this paper, we demonstrate that first-order Markov assumptions limit accurate modeling of NIS spread risks through the global shipping network. In contrast, we show that higher-order patterns overcome this limitation by revealing indirect pathways of NIS transfer. We accomplish this by developing Species Flow Higher-Order Networks (SF-HON), which we developed independently for ballast and biofouling, for comparison with first-order Markovian models of ballast and biofouling. We evaluated SF-HON predictions using the largest available datasets of invasive species for Europe and the United States. We show that not only does SF-HON yield more accurate NIS spread risk predictions than first-order models and existing higher-order models, but also that there are important differences in NIS spread via the ballast and biofouling vectors. Our work provides information that policymakers can use to develop more efficient and targeted prevention strategies for ship-borne NIS spread management, especially as management of biofouling is of increasing concern.

## 1 Introduction

The subset of non-indigenous species (NIS) that become invasive pose significant and growing threats to ecosystems, human and animal health, infrastructure, the economy, and cultural resources and economic losses, resulting in an estimated annual cost of 1.4 trillion USD globally (of which 120 billion USD is the estimated cost in the United States) [1]. In the Great Lakes alone, the annual cost of ship-borne NIS is estimated as high as 800 million USD [1]. While there are various mechanisms that can lead to the introduction of NIS, this paper focuses on the NIS risks stemming from global commercial shipping, which is responsible for more marine NIS introduction around the world than any other mechanism [2, 3, 4], and whose risk continues to rise with increasing trade and new shipping routes [5].

Prevention of NIS transfer and establishment has been identified as the most effective and economically efficient means to reduce NIS costs [6, 7]. However, prevention requires understanding of and targeting the vectors of NIS introduction. Ship-borne NIS are transported via two main vectors: *ballast water* and *biofouling*. Ballast water management has received significant attention from researchers and policy-makers in recent years, including the International Maritime Organizations International Convention for the Control and Management of Ships’ Ballast Water and Sediments (BWM) and associated national legislation in the US and elsewhere. On the other hand, management of biofouling [8, 9] NIS transfer has received relatively less attention [10]. Not only do the two vectors of marine NIS transport require very different management interventions, but they also present challenges on how to effectively model their respective pathways to more accurately assess risk. Accurate risk assessment is critical to inform a cost-effective and efficient strategy for NIS risk management.

We posit that the NIS spread through the global shipping network is not a simple first-order Markov process as mostly assumed in the literature [11, 12, 13, 14, 15]. Rather, the pathways of NIS introduction exhibit **higher-order pathways of species transfer**, because many ships do not discharge their entire ballast water or release all of their biofouling species in their first port of call. For example, consider the case where a ship visits port *C*, *A*, and then *E*. Suppose the ship takes in ballast water in port *C*, and discharges a portion of water in port *A*, and discharges the rest in port *E*. This means that there is still a considerable risk of species introduction from port *C* to port *E*, in addition to port *A*. Figure 1 (a) left displays an illustration of this situation. The first-order connections may have higher species transfer risk, but a there is a non-trivial possibility of species transfer via either of the mentioned mechanisms from port *C* to port *A*, and *E*. Recurrence of such movement patterns over time results in the formation of indirect pathways for species introduction in the long-term. The above scenario holds for biofouling as well. Species that accumulate on ship body in port *C* can still impose introduction risk to port *E* in addition to port *A*, if they manage to survive along the path *C* → *A* → *E*. Ignoring such dependencies will result in inaccurate estimation of the introduction risk.

**Figure 1:**
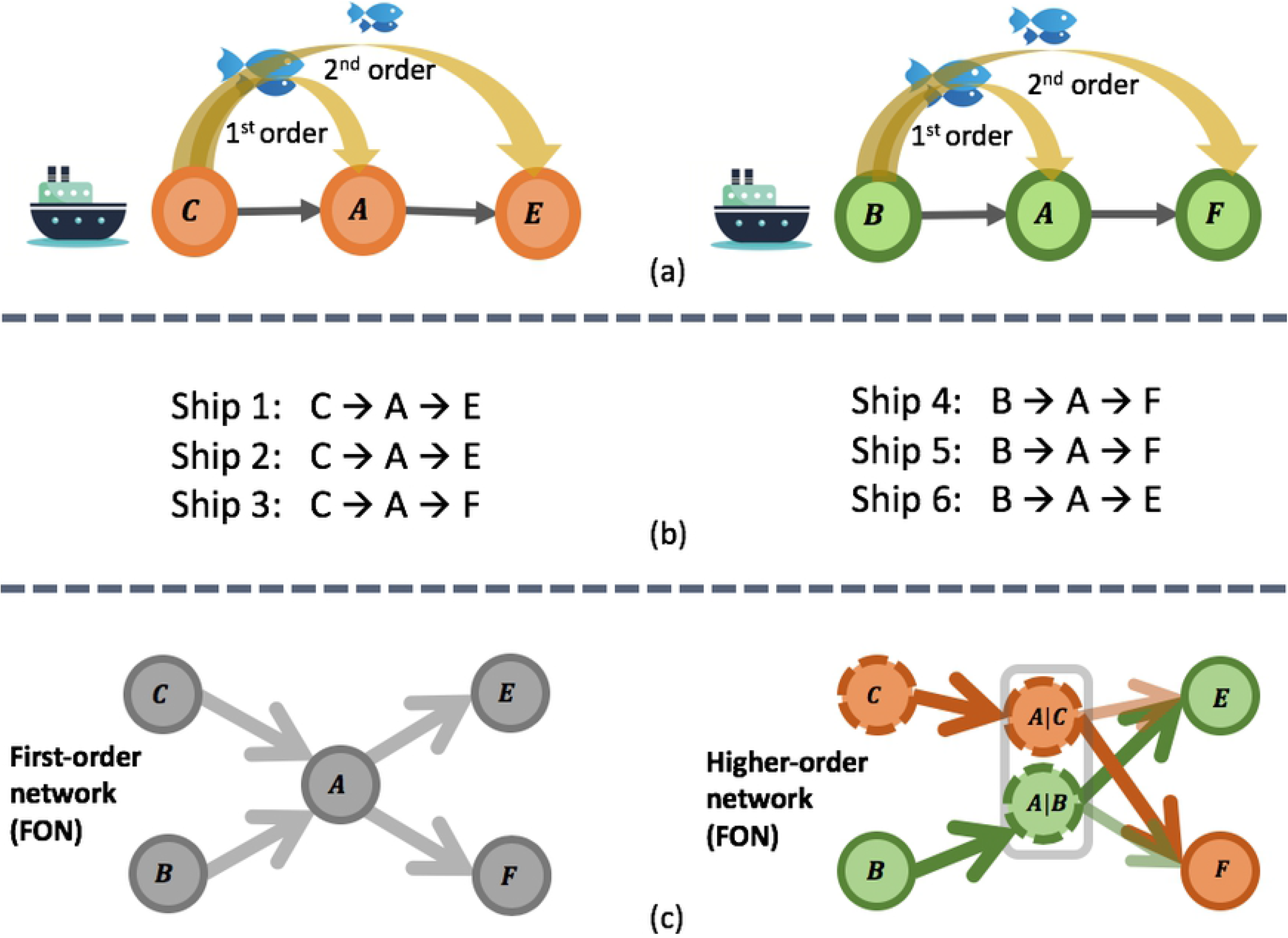
Two different pattern of higher-order species transfer between ports. (b) Out of six ships, ship 1 and 2 follow the pattern in (a) left, and ship 4 and 5 follow the pattern in (a) right. Conventional first order network ((c) left), where organisms have the same probability of being introduced to port *E* and *F* via port *A*. Higher-order network ((c) right) with a second-order dependency example in species transfer. Due to higher-order patterns in ship movements, organisms native to port *C* are more likely to get transferred to port *F* via ships traveling through port *A* and organisms native to port *B* are more likely to get transferred to port *E* via ships traveling through port *A*.

Previous works have recognized the importance of higher-order dependencies in ship-born NIS transfer and some have attempted to incorporate them into their risk models. The authors in [16, 17] propose to include the entire ship trajectory for calculating the NIS risk. However, some connections are more important than the others. For example, only the significant and reoccurring higher-order patterns are likely to result in species introduction at a global scale. As a result, accumulating the risk over the entire trajectory can result in over-estimation of the risk. In [10], the authors consider shipping paths of length five and less to be connected to each other. The problem with this method is that a fixed order is not realistic for all connections: as we explain further in the paper, several longer paths (having up to 15th order of dependencies) could exist in the network. None of the previous models compare higher-order patterns of species introduction through biofouling and ballast discharge. Finally, none of the above models perform a large-scale validation of their model against the first-order models considering the environmental dependencies in higher-order introduction pathways.

### Contributions

Here, we conduct a global comparative study to assess the risk of NIS spread via biofouling and ballast water through the global shipping network. To this end, we develop a species-flow higher-order network (SF-HON) for both introduction vectors by integrating the vessel movement data, environmental data, and bio-geographical data while accounting for all significant higher-order dependencies in the ship movements. We then compare the SF-HON to the first-order network(SF-FON) and another higher-order model in [16] to evaluate predictions of each against NIS introduction datasets from the USA and Europe. This study presents the first large-scale validation of higher-order NIS spread via ballast water and biofouling. Our results can inform policymakers to impose more effective targeted prevention strategies for NIS spread management. Our main contributions include:

- Proposing a global model for biofouling risk by integrating the shipping, environmental and bio-geographical data.
- Presenting a global comparison of the NIS spread risk via ballast water and biofouling. We identify the main differences in network structure, clustering patterns, main pathways, and illustrate the correlation of vessel characteristics and NIS spread risk for each vector.
- Quantifying the impact of incorporating higher-order dependencies into species-flow modeling by comparing the accuracy of SF-HON model against existing higher-order approach and the SF-FON model using two large datasets of NIS from the USA and Europe.
- Illustrating changes in NIS spread risk and SF-HONs characteristics over a span of 15 years of shipping data.

## 2 Materials and methods

### 2.1 Data sources

As illustrated in the block diagram of Figure 2, we incorporated the shipping and environmental data sources into SF-HON. We then used invasion records from *AquaNIS* data and the *USGS* data for validating the effectiveness of SF-HON. All validation datasets are filtered to include records from 1997 to present. We assume it takes one to two decades for spices to be introduced via shipping activity, thrive and reproduce in the new environment to be publicly detected as an invasive species.

**Figure 2:**
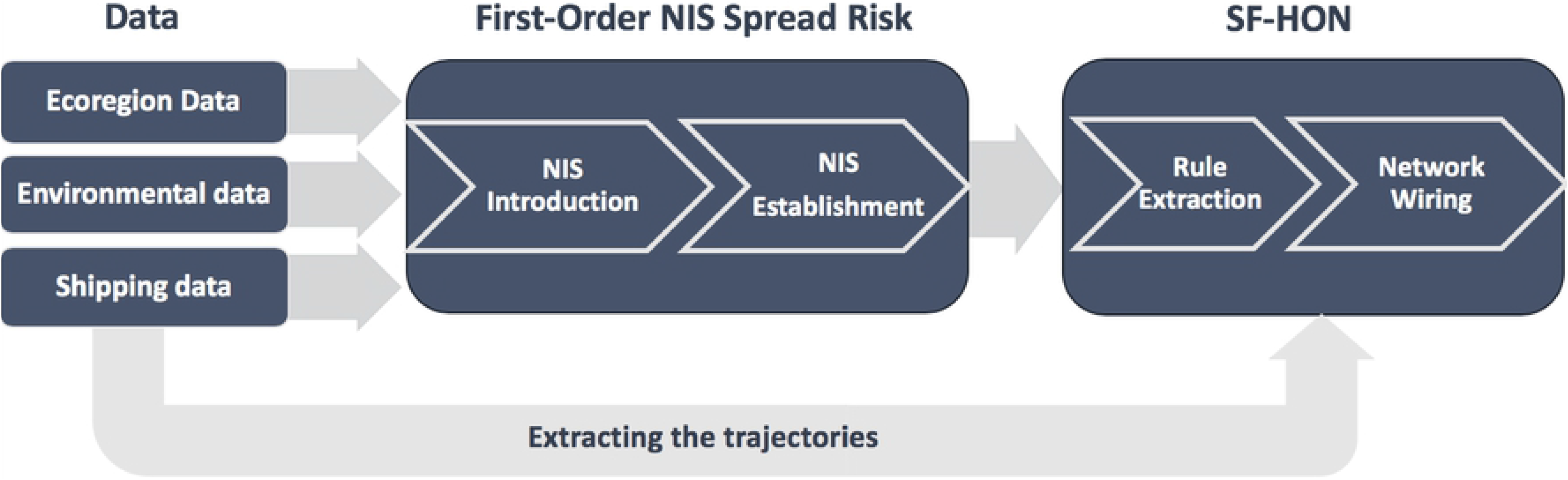
A block diagram of the risk network construction. The ecoregion data, environmental data, and the shipping data are used to compute the first-order (pairwise) NIS spread link between the shipping ports through each vessel introduction vector. The resulting probabilities along with the shipping trajectories is used for constructing the two SF-HONs, ballast SF-HON and biofouling SF-HON.

#### Vessel movement data

We extracted the ship movement patterns using Lloyd’s Maritime Intelligence Unit (LMIU) database. This data contains travel information for vessels such as port ID, sail date and arrival date, along with the vessel meta-data. This information is used to build the first-order and higher-order network to represent the risk of invasion through ballast discharge and biofouling among ports. Our study is based on LMIU data over the most recent year of data available to us starting May 2012 with 1,185,510 individual voyages. However, for section 3.3 we used the entire 15-year period of data starting 1st of May 1997, with 9,482,285 voyages.

#### Ballast discharge data

Provided by the National Ballast Information Clearinghouse (NBIC) [18] contains the date and the discharge volume of all ships visiting U.S. ports from Jan. 2012 to present. We used the NBIC data to estimate an average ballast discharge for ships given their type and size.

#### Environmental data

The temperature and salinity of the shipping ports is extracted from the Global Ports Database [10] and the World Ocean Atlas [19] using the approach in [10]. From 9177 ports in the vessel movement data, 6695 ports with the temperature and salinity information were included.

#### Biogeographical data

Ecoregions are geographical regions of the ocean that share the similar species sets due to shared evolutionary history. We obtained the ecoregion data from Marine Ecoregion of the World (MEOW) [20] to determine the likelihood of a transport species being non-indigenous given the source and destination ecoregion.

#### AquaNIS nonindigenous species data

This data set contains information on NIS introduction histories, recipient regions, taxonomy, biological traits, impacts, and other relevant documented data. This data set is constantly updated with new records and is regarded as best available [21]. The NIS data is available for species introduced to the European countries and is filtered to only include species suspected to have been introduced by ballast or biofouling from 1997 to present.

#### USGS nonindigenous species data

This is the non-indigenous aquatic species database from the United States Geological Survey (USGS). This dataset contains reliable information about the distribution and presence of species across the United States. Most of the species in this dataset are freshwater species. Data fields include the information about the introduced species such as taxonomy, state, drainage unit, freshwater/marine. We extracted the first introductions for each NIS in each state that were potentially transported by ballast water or biofouling and recorded between 1997 to present.

#### National Exotic Marine and Estuarine Species Information System (NEMESIS)

This is the non-indigenous marine and estuarine species database for North America. Taxonomic coverage includes algae and animal species. Data fields include taxonomic descriptions and classification, global distribution, invasion history, ecological and life history traits, and known environmental and economic impacts. We extracted the first introduction for each NIS in each state that was potentially transported by ballast water or biofouling and recorded between 1997 to present.

### 2.2 Risk assessment method

In this section, we explain our calculation of first-order (pairwise) NIS risk, construction of SF-HON and calculation of higher-order risks, network analysis methods used, and the evaluation method.

In order to calculate NIS risk for the species flow first-order networks (SF-FONs), pairwise NIS risks were summed for each port pair. For species flow higher-order networks (SF-HONs), we first identified significant higher-order dependencies or “rules” (Figure 1 using a technique developed by [22] and then re-constructed the network accordingly. The higher-order NIS risk is calculated using the SF-HON.

In addition to comparing network structure and clustering patterns between ballast and biofouling SF-FONs and SF-HONs, we also tested how well each model predicted recent NIS introduction data from Europe (*AquaNIS*) and the USA (*USGS*). Below we describe each step in more detail.

#### 2.2.1 The NIS spread

As illustrated in the block diagram of Figure 2, we incorporated the shipping, environmental, and biogeographical data sources to estimate NIS spread risk for each ship voyage as the product of three independent probabilities: probability of introduction to port, probability of being non-indigenous to the destination port and probability of establishment (survival and reproduction). NIS introduction risks for ballast water and biofouling were calculated using a different formula, yielding separate NIS spread risks for these two vectors. For both introduction mechanisms we define the invasion risk between port *i* and *j* to be the product of the three probabilities:

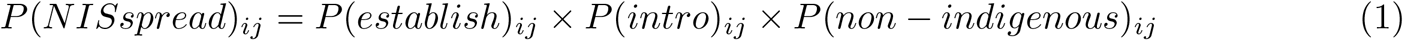

##### The probability of being non-indigenous

We identified ports pairs that may contain non-indigenous species based on the ecoregion data, to account for the fact that many wide-spread species are found in more than one ecoregion [23], [20] and many are able to disperse to neighboring ecoregions naturally. As a result, neighboring ecosystems are likely to share more species that are considered native [23].

Therefore, if the source and destination ports belong to the same or neighboring ecoregions, the probability that they introduce a non-indigenous species to each other is 0; otherwise, this probability is set to 1.

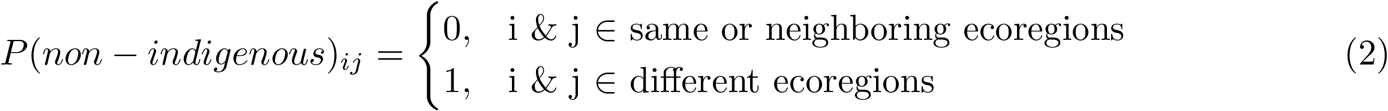

##### The probability of establishment

This probability is calculated based on the environmental similarity of the give ports. We adapt the state-of-the-art formulation and parameters for this part [16]: This probability is modeled as a Gaussian distribution of the temperature difference Δ*T*_*ij*_ and salinity difference Δ*S*_*ij*_ of the source and destination ports, normalized by their standard deviation, *δ*_*T*_ and *δ*_*S*_,

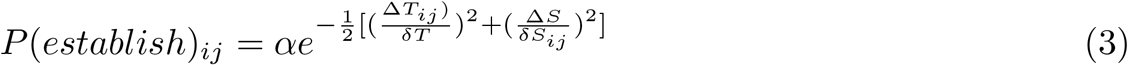

in which *α* = 0.00015, *δ*_*T*_ = 2°*C* and *δ*_*S*_ = 10 *ppt* are chosen based on [16].

##### The introduction probability via biofouling

Fouling risk is calculated based on two main factors: species accumulation and species survival during the trip.

We estimate the relative biofouling accumulation on a ship based on two parameters: the duration of stay at the source port *i*, *d*_*i*_, and the antifouling practice based on the ship type *A*^(*t*)^. The relationship between biofouling accumulation (measured as the proportion of maximum species richness) and duration in port *i* (measured in days) was derived from data taken from four studies which had tracked biofouling community accumulation over time at multiple latitudes [24], [25], [26], [27]. The relationship was modeled as a 3rd order polynomial, with accumulation at locations in tropical and subtropical latitudes (equator +/− 35 degrees latitude) and temperate latitudes having different patterns (figure S1, supplementary materials). The antifouling parameter *A*^(*t*)^ refers to the proportion of ships of a given type without an operational antifouling system. Estimates for each ship type were obtained from a survey of commercial ships in California [28] as follows: Container Ships: 0.19, Automobile Carriers: 0.20, Tankers: 0.30, Passenger Ships: 0.31, Bulk Carriers: 0.42, and General ships: 0.53, and all other commercial ships: 0.60.

The second main factor in the introduction risk via biofouling is the survival probability of species during the voyage, which is known to decrease with increasing voyage velocity *v*_*ij*_. Using experimental data from [29], we fit an exponential decay function to estimate survival probability as a function of *v*_*ij*_ (figure S2, supplementary materials).

Considering both species accumulation and survival probability, we estimate biofouling introduction risk for each voyage between port *i* and *j* as:

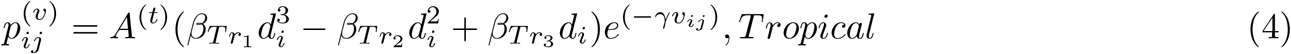

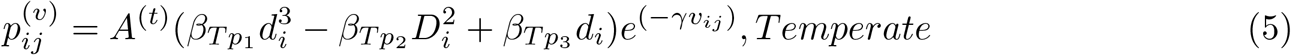

Where *v*_*ij*_ is the average voyage velocity from source port *i* to destination port *j* measured in kilometers per day. *γ* = 0.008 and *β*_*Tr*1_ = 1.29 × 10^−7^, *β*_*Tr*2_ = 8.316 × 10^−5^, *β*_*Tr*3_ = 0.0149 are the tropical coefficients and *β*_*Tp*1_ = 1.4 × 10^−9^, *β*_*Tp*2_ = 1.6566 × 10^−5^, *β*_*Tp*3_ = 5.193 × 10^−3^ are the temperate coefficients (refer to section 1 in supplementary materials for more details on model development and calculation details).

##### The introduction probability via ballast discharge

Our method for estimating introduction probability based on ballast discharge slightly alters the equation used by [16]. Let *v* be a ship traveling from port *i* to *j*, during the time 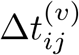. Species in the ship ballast water may die at a daily mortality rate of *μ*. Let 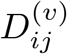, *ρ*^(*v*)^ [0, 1], and λ be the amount of ballast water discharged at the destination, the efficacy of ballast water management for the route, and the species introduction potential per volume of discharge. Then, the probability of vessel *v* introducing species from *n*_*i*_ to *n*_*j*_ is given by [16]:

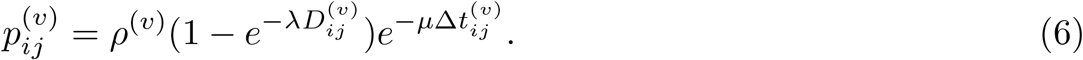

The amount of ballast water 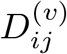 is estimated based on ship type and ship gross weight tonnage using the method proposed in [14]. The trip duration 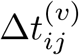 is extracted from Lloyd’s data set. *μ* = 0.02 and λ = 3.22 × 10^−6^ are chosen based on [14].

For both introduction mechanisms, the total probability of introduction is the aggregated risk of all ships traveling from *i* to *j*:

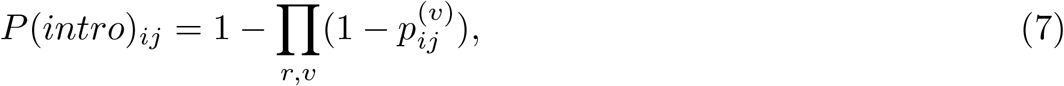

where the product is taken over all routes *r* such that a ship *v* travels from port *i* to *j*.

### 2.3 SF-HON: Higher-order network of species flow risk

We calculate the NIS spread risk for every pair of ports under ballast discharge and biofouling mechanisms using the model defined in Section 2.2. At this point, we could conventionally model this data into a network structure where nodes represent the shipping ports and edges correspond to the first-order port risks. However, this simple interpretation is an over-simplification of the complex system of the global shipping traffic. Studies show that a ships’ next ports to visit can depend on ships’ previously visited ports [22]. As illustrated in Figure 1(b), ships coming to port *A* from port *C* are more likely to visit port *E* thank *F*, and ships coming to port *A* from port *B* are more likely to visit port *F* , than port *E*. These patterns indicate a second-order dependency, which implies that species native to port C has a higher introduction probability to port E (via port A) and species native to port B have a higher introduction probability to port F (via port A, as illustrated in Figure 1(a)). However, if first-order movement pattern is assumed (Figure 1(c) left), the ship has equal probabilities of visiting either port *E* or *F*. Considering only first-order interactions does not capture such introduction probabilities. Therefore, a more accurate way of wiring the shipping ports is required to account for these hidden patterns. The Higher-Order Network (HON) representation is shown in figure 1(c) right, in which probability of going to port *E* and *F* from port *A* varies given the port visited before port *A* (i.e., probability of going from *A*|*C* to *F* is higher than probability of going from *A*|*C* to *E*). Such higher-order patterns are also very important in the environmental conditions of the ports. For example, species that are intolerant to cold temperature will probably die in port *P*_2_ if the ship travels a path of *P*_1_(*tropical*) → *P*_2_(*arctic*) → *P*_3_(*tropical*). Therefore, considering only first-order interaction of *P*_1_(*tropical*) and *P*_3_(*tropical*) does not represent the intermediate port *P*_2_(*arctic*) and its environmental conditions.

Xu et al., [22] proposed a framework to model such higher-order movement patterns in a network structure. Here we adapt this framework to assess the higher-order NIS spread risk. From a total of 1,185,510 voyages during 2012-2013 (latest available year of shipping data), we extracted 29,788 and 49,895 unique first-order pathways (pairwise connections without considering intermediate ports). For each pathway, we calculate the pairwise biofouling and ballast discharge risks between source and destination ports based on the available data (ecoregion data, environmental data, and the shipping data). We also construct the vessel trajectories by listing the sequence of ports visited by each vessel during a one year period. Then, we combine the pairwise risks together with the vessel trajectories to determine which indirect shipping routes exhibit higher-order species introduction patterns and produce the Species Flow Higher-Order Network (SF-HON). As a result, SF-HON contains the higher-order pathways of NIS transfer in the network structure. A block-diagram of the work-flow is illustrated in figure 2. We use the vessel trajectories and the NIS spread risk of the paths as the input for SF-HON. We then feed these probabilities to the Rule Extraction step to identify significant higher-order dependencies from the sequence of trajectories. The generated rules will be used to generate the SF-HON in the Network Wiring step.

#### Rule extraction

In the rule extraction step, the objective is to identify the significant higher-order introduction pathways by finding the correct order of dependency in the data. We use the similar rationale as [22]: Given a pathway of order 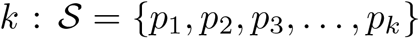, We can calculate the corresponding NIS spread probability of the path using the below formula:

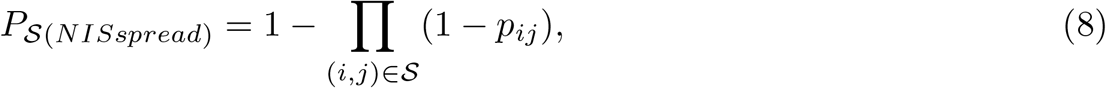

In order to identify the correct order of dependency based on the shipping trajectories, using a similar approach as [22], we check whether including a previous step *p*_0_ and extending 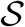 to 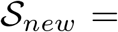 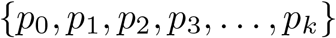 (of order *k*_*new*_ = *k* + 1, with NIS spread probability of 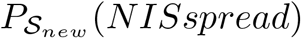) will significantly change the NIS spread probability of the path. If so, order *k*_*new*_ is assumed as the new order of dependency, and 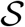 will be extended to 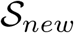. The resulting rule is 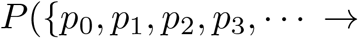 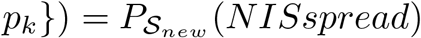. The above method only accepts rules that are significant and have occurred sufficiently enough times. Therefore, random patterns will not appear as a dependency rule. Furthermore, our method allows for variable order of dependencies for different paths.

#### Network Wiring

In the network wiring step, the extracted rules are converted to corresponding nodes and edges in the SF-HON. For example, given a discovered rule *P* ({*p*_0_, *p*_1_, *p*_2_ → *p*_3_}) = 0.5 we define the edge *p*_2_|*p*_0_, *p*_1_ → *p*_3_ with the weight 0.5. This indicates that the probability of NIS spread from *p*_2_ to *p*_3_ for ships who have visited ports *p*_0_ and then *p*_1_ before coming to *p*_2_ is 0.5. The node *p*_2_|*p*_0_, *p*_1_ is a third-order node, and *p*_3_ is a first-order node. The resulting network is called SF-HON in which a node can represent a sequence of ports, and thus several nodes in SF-HON can map to a single physical port (for example, *p*_2_|*p*_0_, *p*_1_ and *p*_2_|*p*_1_ both represent the physical port *p*_2_). We construct the SF-HON for the two vessel introduction vectors and define them as *ballast SF-HON* and *biofouling SF-HON*. We perform all the further analysis based on these two SF-HONs.

### 2.3.1 Network Clustering Analysis

Clustering allows us to obtain a large-scale view of NIS spread and identify the groups of ports which have a higher (or lower) probability of introducing species to each other. Ports within the same cluster are relatively more connected and have a stronger chance of species spread. Note that, using the higher-order approach, each physical port can belong to several clusters, because its corresponding higher-order nodes belong to different clusters. For example, *p*_2_|*p*_0_, *p*_1_ belongs to cluster 1 and *p*_2_|*p*_1_ belongs to cluster 2, while they both represent the physical node *p*_2_.

We used *Infomap* [30] as a network clustering method on the two risk networks. The basic principle of Infomap clustering is to find groups of nodes among which information flows quickly and easily and can be aggregated as a separate cluster. This clustering method is suitable for extracting modules of species flow since it identifies clusters by optimizing the entropy corresponding to intra-cluster and inter-clusters using a recursive random-walk method, and random walks are the most similar to the species flow pattern.

### 2.3.2 Evaluation metrics

In this work, our goal is to demonstrate that SF-HON yields more accurate predictions for NIS spread risk. To verify that, we compare the results from SF-HONs with the conventional SF-FON and another higher-order model proposed by [16]. In [16], the authors include the entire shipping trajectory for calculating the NIS risk, without any further constraints on the significance of the higher-order patterns. We refer to the baselines using this method as *Ballast Full-Traj* and *Biofouling Full-Traj*.

We evaluate the SF-HON predictions against SF-FON and Full-Traj baselines using one dataset for European countries (the AquaNIS data) and two datasets for United States (the USGS and NEMESIS data). We filtered all three datasets to include only species suspected to have been introduced by ballast or biofouling from 1997 (the first year of our shipping data) to present. For the US, we obtained a total of 251 species introduction records from USGS and 250 species introduction records from NEMESIS. We define the *NIS introductions* to be the normalized count (between 0 and 1) of the first introduction for each NIS reported from each state. For AquaNIs data, we extracted a total of 815 introductions for 20 countries, and define the *NIS introductions* to be the normalized count (between 0 and 1) of the first introduction for each NIS reported from each country. Normalization is done so that we can compare our results across different models and different datasets.

We extracted the NIS spread risk scores from ballast SF-HON and biofouling SF-HON along with the SF-FON and Full-Traj models by averaging the obtained NIS spread risks over all available ports corresponding to each state (and each country, in case of Europe). We then calculated one score for each network model by averaging over all the ports within a State and country, and scaled the final scores over the available states and countries. We define the MSE for each model by calculating the square difference between the *NIS introductions* and the scaled NIS spread risk obtained from the network models. We calculate MSE for each state and each country and report the average. (for detailed results on each state and country, refer to figure S6 in supplementary materials)

## 3 Results

Using the model defined in Section 2.2, we obtain two NIS spread risks corresponding to ballast discharge and biofouling introduction mechanisms for every pair of ports. We first conventionally modeled this data into a network structure where nodes represent shipping ports and edges correspond to the first-order port risks (ballast SF-FON and biofouling SF-FON). We then went further and significant higher-order patterns of ship movements as described in [22] to create the ballast SF-HON and biofouling SF-HON. Below we provide a comparison of the SF-HONs and SF-FONs in terms of basic network characteristics. We then compare ballast and biofouling SF-HONs in terms of NIS flow across major biogeographic realms, the evolution of global NIS flow patterns from 1997-2012, and global clustering patterns. We end by evaluating SF-HON, SF-FON and an existing higher-order model [16] against large NIS introduction datasets from the USA and Europe. We did not include a higher-order network analysis using [16], since higher-order pathways in this method depend on the length of ship trajectories and not significant reoccurring patterns. As a result, higher-order patterns in this model do not contain any additional signals. Furthermore, constructing a higher-order network based this approach results in a very sparse network which does not allow for network analysis methods like clustering.

### 3.1 Network characteristics

We first evaluate the basic network properties of the two SF-HON (ballast and biofouling) and compare them with their first-order counterparts (SF-FON). Our analyses indicate that ballast water transport creates significantly more higher-order nodes and NIS spread pathways than does biofouling transport.

If no higher-order dependencies exist in the risk network, the SF-HON and SF-FON would be similar. However, this is not the case in ballast SF-HON. Table 1 summarizes the properties of the two risk networks. From this table, we can observe that the ballast SF-HON expands significantly (relative to the SF-FON) due to the inclusion of significant higher-order patterns: the number of nodes and edges grow 37.56 and 5.08 times larger than SF-FON, respectively. This increase is not as large in the biofouling SF-HON: number of nodes and edges in SF-HON become 9.39 and 3.13 more than SF-FON, respectively. This indicates that there are significantly more higher-order pathways of species transfer via ballast discharge. For example, species are likely to get introduced from *Singapore* to *Busan* via path: *Singapore* → *Shanghai* → *Tokyo* → *Busan*, while in biofouling SF-HON species mostly get introduced via first-order paths, (e.g. *Singapore* → Shanghai, or *Tokyo* → *Busan*). For more detailed compassion of network degree distribution and clustering coefficient, refer to figure S3 in supplementary materials.

**Table 1:**
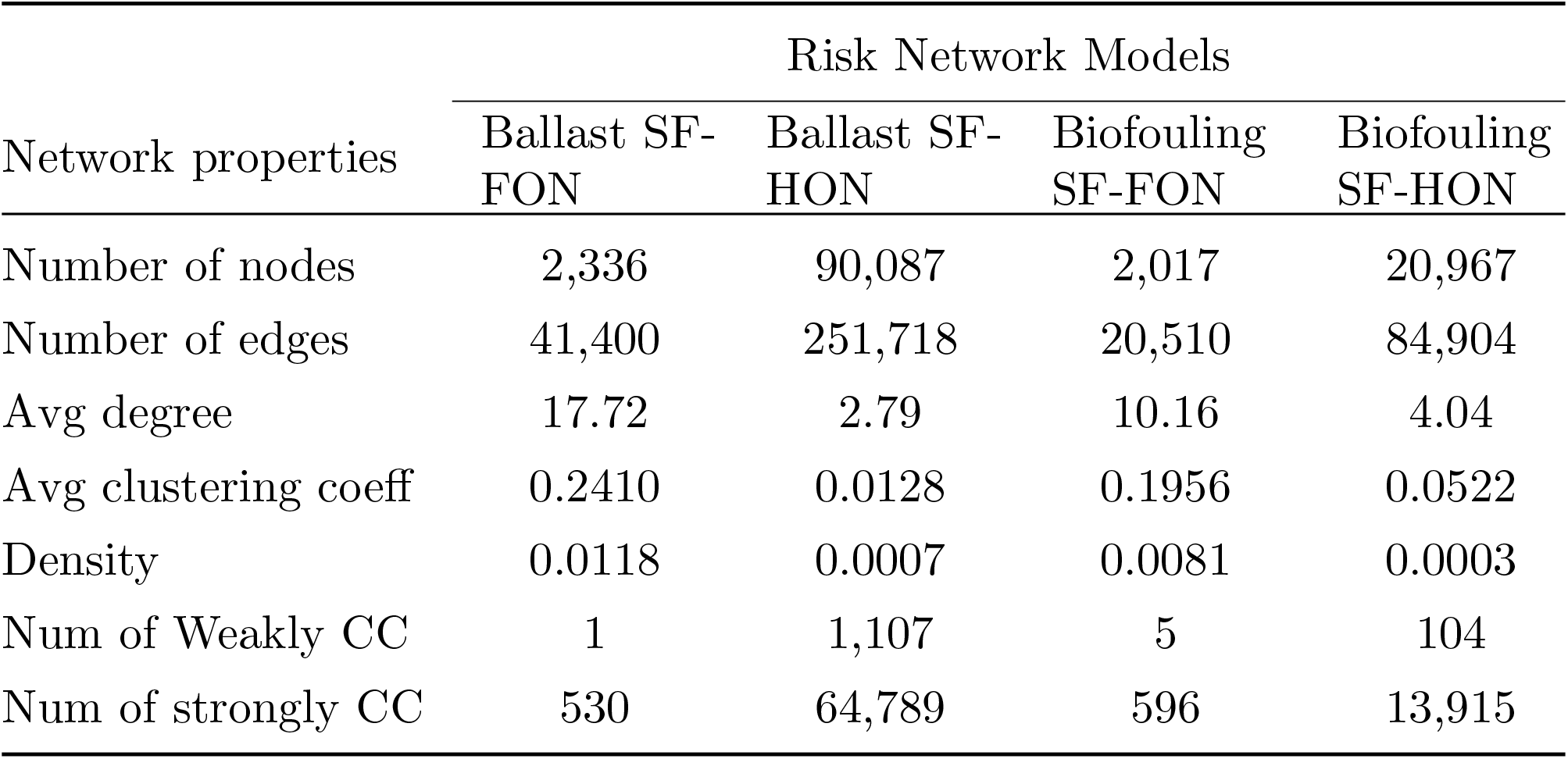
Comparison of basic properties of the four networks

**Table 2:**
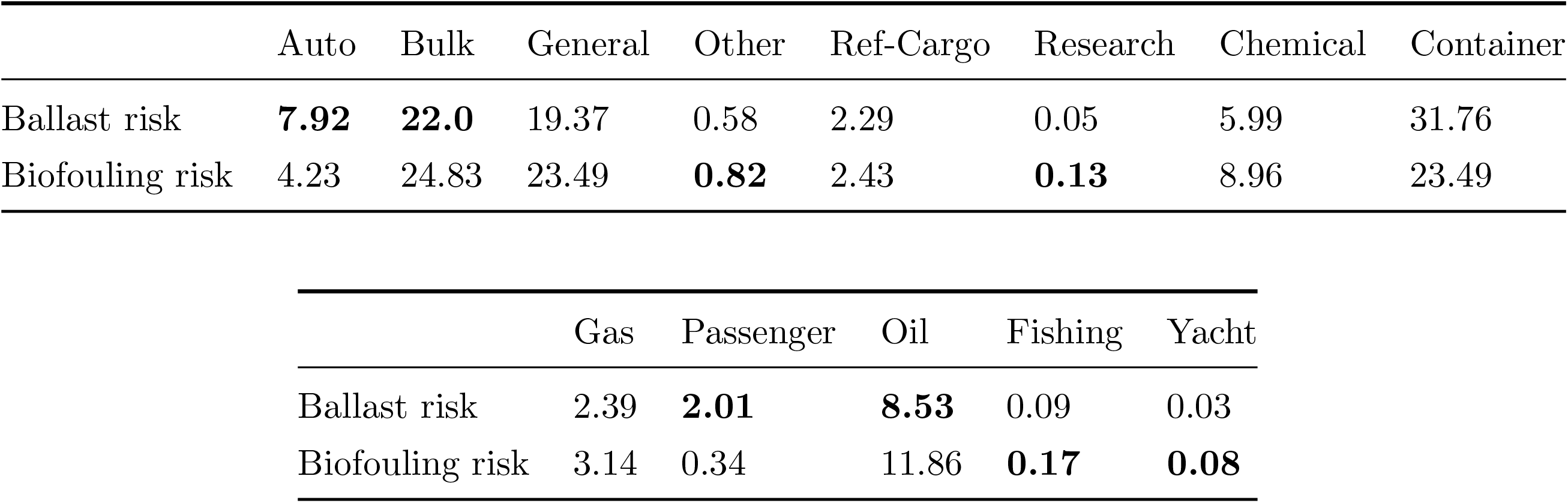
Percentage of each vessel type in the data for calculation of ballast and biofouling risk.

|  | Auto | Bulk | General | Other | Ref-Cargo | Research | Chemical | Container |
| --- | --- | --- | --- | --- | --- | --- | --- | --- |
| Ballast risk | **7.92** | **22.0** | 19.37 | 0.58 | 2.29 | 0.05 | 5.99 | 31.76 |
| Biofouling risk | 4.23 | 24.83 | 23.49 | **0.82** | 2.43 | **0.13** | 8.96 | 23.49 |

|  | Gas | Passenger | Oil | Fishing | Yacht |
| --- | --- | --- | --- | --- | --- |
| Ballast risk | 2.39 | **2.01** | **8.53** | 0.09 | 0.03 |
| Biofouling risk | 3.14 | 0.34 | 11.86 | **0.17** | **0.08** |

These differences between the ballast and biofouling SF-HONs have implications for global NIS spread patterns. To compare large-scale patterns of species flow in both SF-HONs, we calculated the NIS spread risks between realms (large-scale biogeographical regions) by averaging over all the ports within a realm. We visualize results as two heatmaps for ballast SF-HON (Figure 3 (a)) and biofouling SF-HON (Figure 3 (b)). We notice that in ballast SF-HON, realms are more likely to introduce species to each other, since the most high-risk connections in ballast SF-HON are intra-realm connections (Figure 3 (b)), while in in ballast SF-HON several inter-realm connections exists. The main high-risk inter-realm connections in biofouling SF-HON are *Temperate Northern Pacific* → *Tropical Atlantic, Tropical Atlantic* → *Western Indo-Pacific, Tropical Eastern Pacific* → *Eastern Indo-Pacific, Western Indo-Pacific* → *Central Indo-Pacific,* and *Arctic* → *Temperate Northern Atlantic* (Figure 3 (b)). This provides a high-level information of the inter-realm NIS spread risk for global management plans. For a breakdown of dependency orders across major realms refer to figure S4.

**Figure 3:**
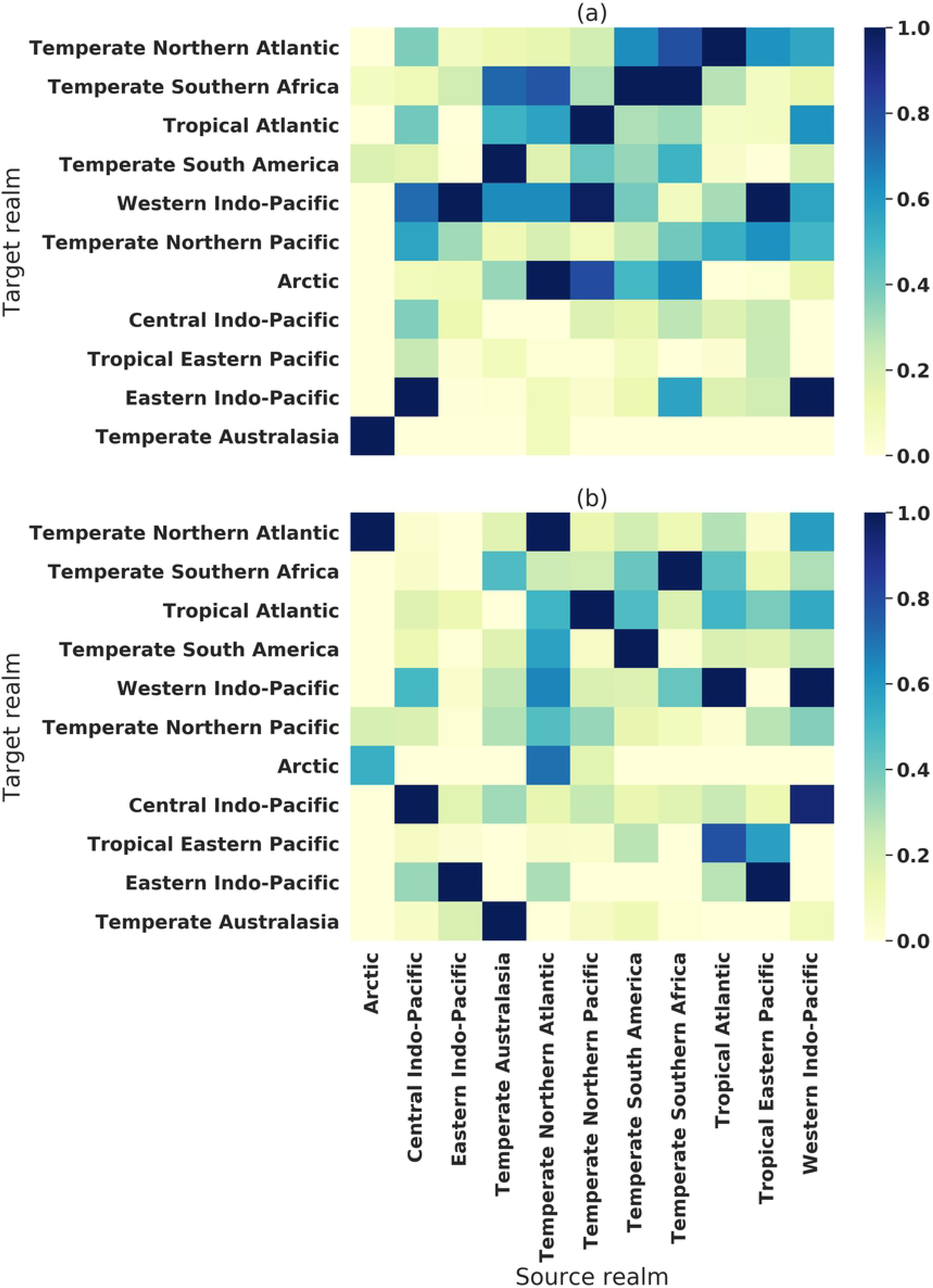
heat-map of higher-order species flow resulting from ballast water across major realms. The y-axis indicates target realms and the x-axis are the source realms (x-axis is shared). The risks are normalized to be in the range of [0,1]. The intra-realm risk is overall higher in ballast SF-HON (a). Eastern-Indo Pacific has the highest risk of NIS introduction to Temp Southern Africa in ballast SF-HON (a), while in biofouling SF-HON the highest target realm for Eastern-Indo Pacific is Central Indo-Pacific (b). Temperate Northern Atlantic has a significantly high risk of introduction to Eastern-Indo Pacific in biofouling SF-HON (b).

### 3.2 Clustering of ballast and biofouling SF-HONs

Clustering allows us to identify partitions of the networks in which ports are tightly coupled by species flow. It also provides a high-level view of interactions of the clusters. Major differences of clustering exist between the ballast and biofouling SF-HONs in terms of geographical organization of clusters, cluster overlap, and distribution of the ports within each cluster.

Even though the same shipping data is used in both networks, Figure 4 (a) shows that small, overlapping, and globally connected clusters predominate in the ballast SF-HON (Figure 4 (a)) while larger, more isolated clusters characterize the biofouling SF-HON (Figure 4 (b)). This highlights the importance of management interventions that could target inter-cluster connections in biofouling SF-HON.

**Figure 4:** The diverse higher order clustering of shipping ports based on the ballast discharge risks (a) and higher-order clustering of biofouling risks (b). Different colors in each pie charts for ports corresponds to different clusters. The pie chart size indicates the relative NIS spread risk for the port. Ports with the highest risk are labeled by name.

Furthermore, figure 4 shows that ballast SF-HON contains significantly more higher-order ports than biofouling SF-HON. Specifically, only 8.64% of the ports in the biofouling SF-HON belong to more than one cluster, while 65.14% of the ports in the ballast SF-HON belong to multiple clusters, which implies that they are more likely to introduce NIS to each other. *This indicates that species spread through ballast water results in significantly more higher-order ports and more clusters.* Overall, the European cluster, the Southeast Asian cluster, and the Indian cluster contain the largest portion of higher-order nodes in the ballast SF-HON. In biofouling SF-HON, only a few clusters contain significant higher-order nodes, namely, the Southeast Asian cluster, the Indian cluster, and the North African cluster. Ports with highest NIS spread risk and the number of clusters that they belong to are also different between the two networks (Table S1, supplementary materials). For example, only 5 ports rank among the top 12 high-risk ports in both the ballast SF-HON and biofouling SF-HON. Additionally, the top 12 high-risk ports in the ballast SF-HON belong to 59-191 clusters, while those in the biofouling SF-HON belong in only 13-80 clusters. Table S1 lists the highest risk ports for ballast and for biofouling.

In summary, we can conclude that clusters in the biofouling SF-HON are more geographically localized with fewer inter-cluster connections. Therefore, targeting the inter-cluster links are a better strategy for preventing biofouling NIS spread from one cluster to another (e.g., [14]). Ballast clusters, on the other hand, have a high inter-cluster connection, and they are less affected by geographical proximity and more influenced by ship traffic, requiring both local and global prevention strategies.

### 3.3 Evolution of SF-HONs and NIS spread risk

In this section, we include the entire shipping data available to us (1997-2012) to analyze changes in network structure and the NIS spread risk for ballast and biofouling SF-HONs.

In both networks, the average NIS spread risk is highly correlated with the number of higher order nodes in the network over time (Pearson correlation value: 0.8362 (*p*-value < 10^−5^)). This indicates that higher-order nodes generally have a higher risk of NIS spread. Large changes in risk occurred over time. In 2008 the number of higher-order nodes and average risk crashed in the ballast SF-HON while they both reached their peak in the biofouling SF-HON. (Figure 5).

**Figure 5:**
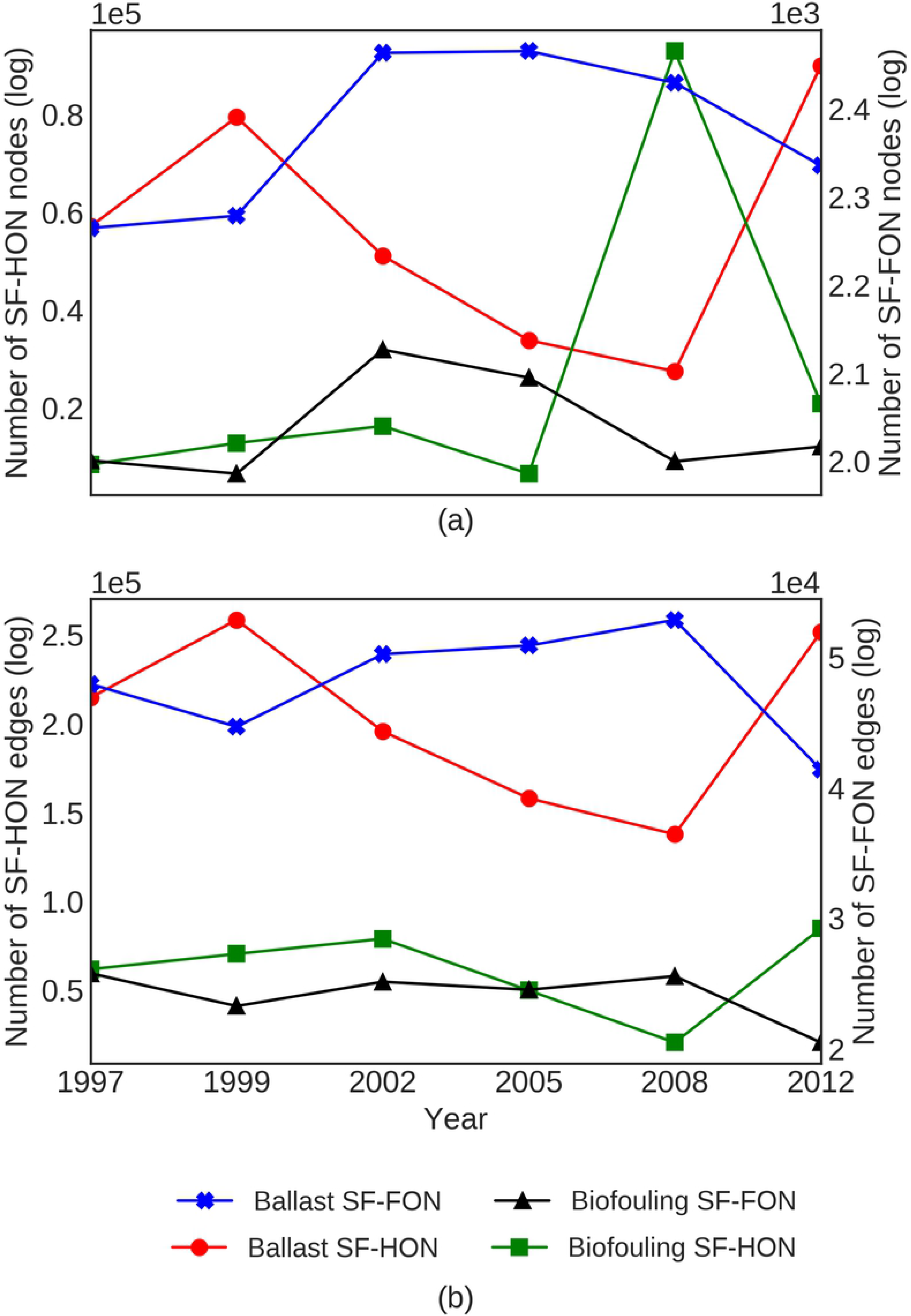
Variation of number of higher-order nodes and average NIS spread risk for (a) ballast SF-HON and (b) biofouling SF-HON. The average risk in both cases is normalized for easier comparison. In 2008 the average risk and number of higher-order nodes in ballast SF-HON drops, while they both increase significantly in biofouling SF-HON.

We believe that this pattern is a result of the global recession (2007-2009), which caused a significant reduction in trade and shipping activities. Specifically, during 2008 we detected increases in average voyage distances (1.89 times more than the average), voyage duration (1.39 times longer than average), and ship stay duration at ports (4.84 times more than the average), and decreases in the number of voyage per port (59.57% less than the average), indicating lower shipping activity. As the ships spend more time at the ports, more species accumulate on the ship hull, causing the peak in biofouling risk (more details in section 3.5 for a comparison of biofouling risk and duration of stay at ports for different ship types). On the other hand, ballast risk drops with increased voyage duration and voyage distance, corresponding to longer connections. We conclude that longer connections which have smaller ballast introduction risks because of increasing mortality rates remained consistent during the recession. Overall, changes in the shipping network over time resulted in dramatically different spread risks for the ballast and biofouling SF-HONs, highlighting the need to consider these two transport mechanisms separately.

We further compared the changes in the number of nodes and edges over time on both SF-HONs and SF-FONs. Figure 6 shows the results. We notice that for both vectors, SF-HONs show significantly larger changes in both number of nodes (Figure 6 (a)) and edges (Figure 6 (b)) over time. In particular, SF-FONs structure does not show a significant change as a result of a major shift in the global trade patterns in 2008.

**Figure 6:**
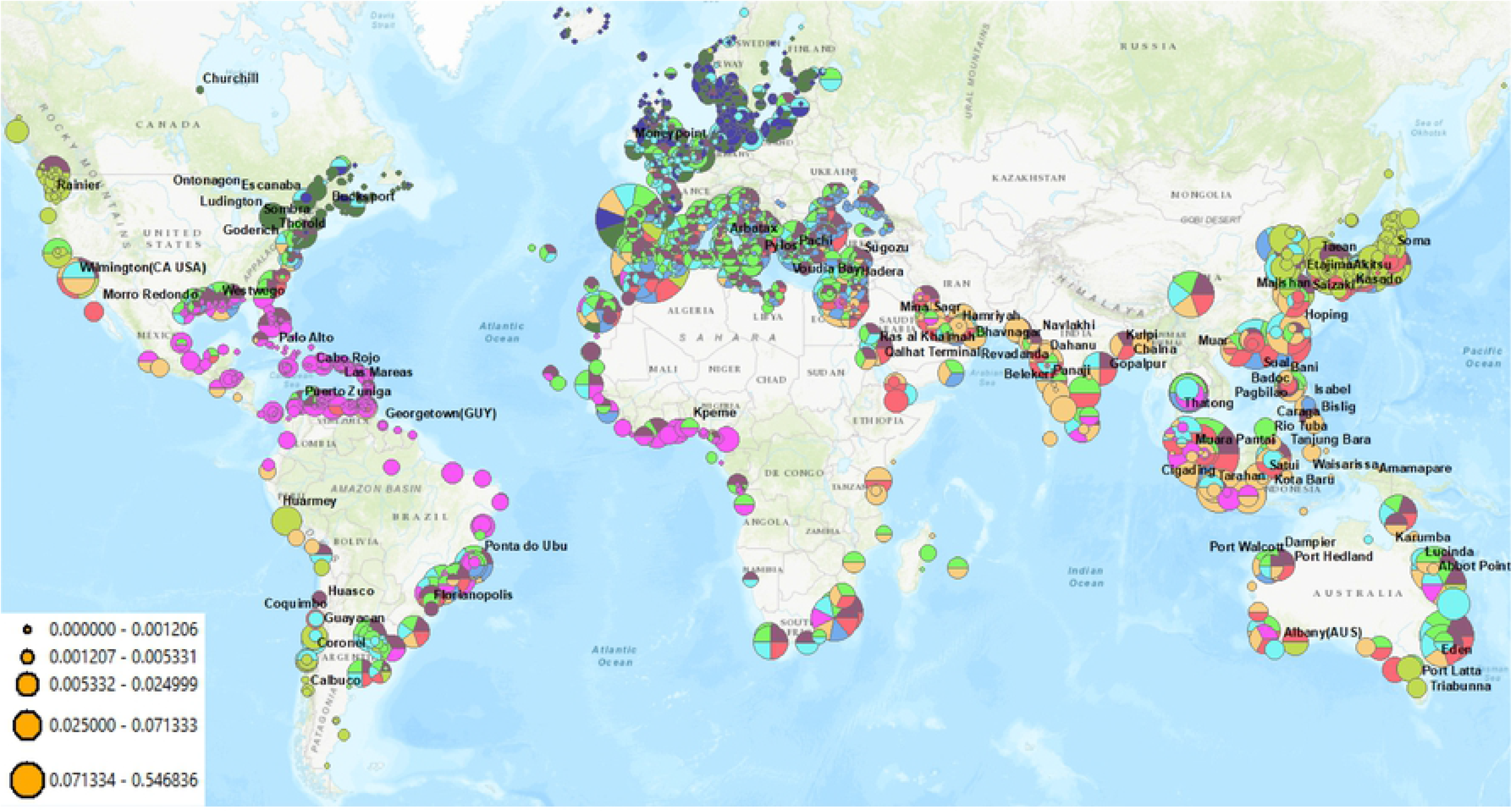
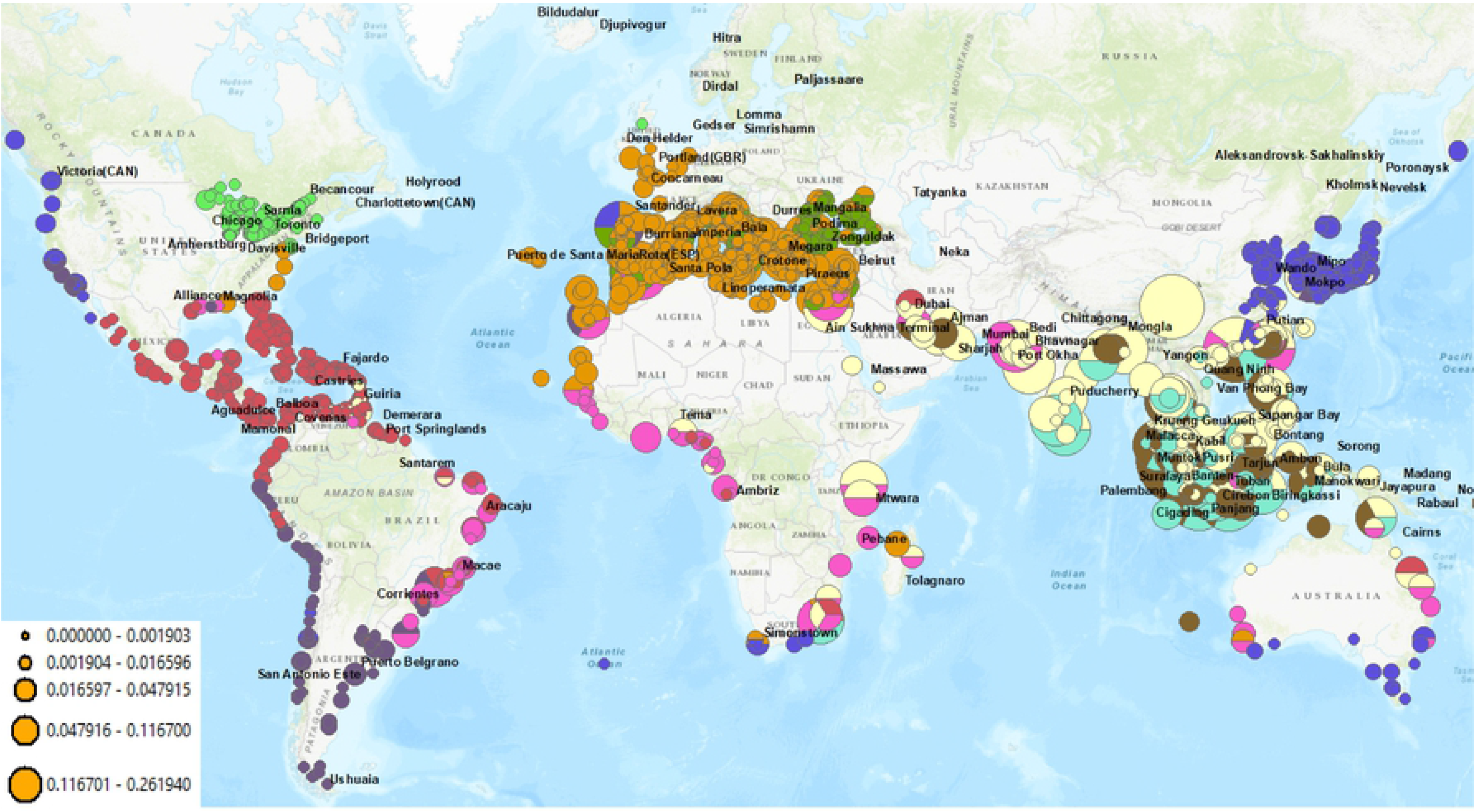
SF-HONs show significantly larger changes in both number of nodes (a) and edges (b). SF-FONs structure does not show a significant change as a result of a major change in the global trade patterns in 2008.

Figure 6 (a) shows that the number of nodes in ballast and biofouling SF-FONs change similarly over time. However, the number of nodes in ballast SF-HON and biofouling SF-HON change in opposing directions. Moreover, Figure 6 (b) shows that the number of edges in SF-HONs change in the opposite direction of SF-FONs. Both observations demonstrate the intrinsic differences in SF-HON and SF-FON as a result of higher-order patterns. A particularly important change point is the significant drop in the number of SF-HON edges in 2008, as a result of the decrease in the shipping activity. We notice that while the number of higher-order ports (and biofouling NIS risk) in biofouling SF-HON increased in 2008, the number of higher-order edges dropped (both in biofouling SF-HON and ballast SF-HON). Despite the drop in the number of edges, biofouling risk increased while ballast water risk decreased in 2008 (Figure 5). This observation shows that the decrease in the shipping frequency has less impact on biofouling risk compared to ballast discharge risk.

### 3.4 Evaluation of SF-HON and SF-FON Predictions

We posit that SF-HONs are a better representation of the species flow patterns as they make more accurate NIS spread predictions than SF-FONs (Figure 7). We compare SF-HON predictions against those of SF-FON and Full-Traj baselines. For each model, we calculated the MSE (Mean Square Error) with respect to previously reported NIS introductions to the USA and Europe (Figure 7; See methods).

**Figure 7:**
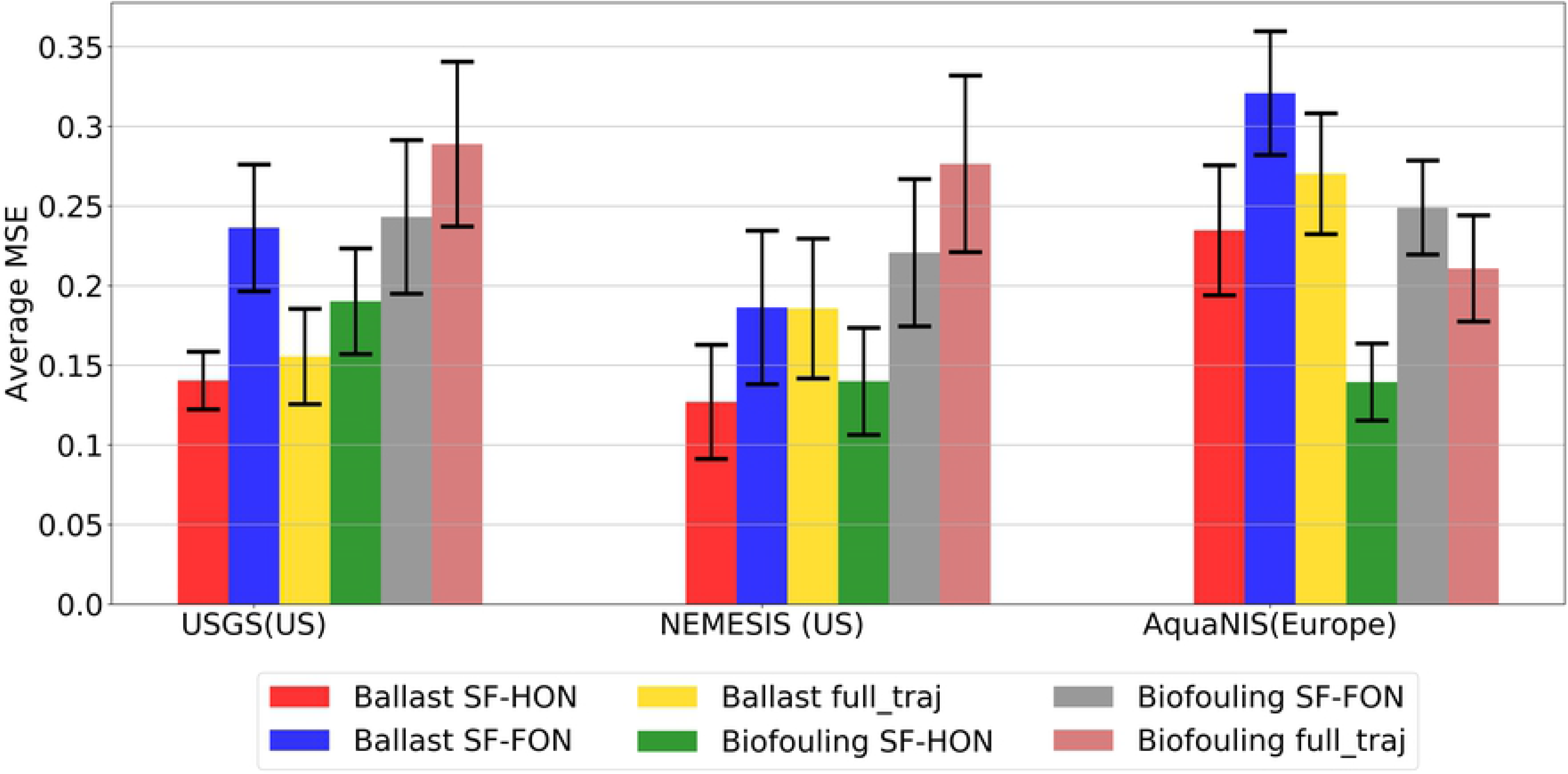
Validation results on the three data sets available. Comparison of the SF-HONs versus SF-FONs and Full-Traj models shows that SF-HONs have generally ballast SF-HON yields more accurate results on the USGS and NEMESIS datasets, while biofouling SF-HON yields more accurate results on the AquaNIS data.

In all datasets, the SF-FONs have higher average MSE. Ballast SF-HON yields the most accurate result for the USGS and NEMESIS dataset, while in the European AquaNIS dataset the biofouling SF-HON performs best. This discrepancy could be due to the fact that the USGS dataset consisted mostly of freshwater NIS records, and ballast water introduction is more likely from international voyages that must cross marine seas where high velocity would kill most biofouling NIS. In the case of Europe, however, smaller voyage distance may make biofouling a more important vector.

We notice that including the entire trajectory in Full-Traj models does not yield more accurate predictions. We investigated this pattern further by looking at individual country/state predictions than SF-HONs. We realized that both Full-Traj models over-estimate the risk in 76.34% of all cases. The first-order models, however, are under-estimating the risk in 79.92% of all cases. We believe the reason is that including the entire trajectory for risk calculation causes the risk from longer pathways to increase the final risk, resulting in over-estimation of the NIS risk. First-order models, on the other hand, do not consider any pathways beyond first-step connections, ignoring the significant higher-order connections, which result in under-estimation of the risk in most cases. SF-HON models provide the most accurate results by considering the significant higher-order pathways. We performed the same analysis over other years as well. SF-HONs consistently perform better than both first-order and Full-Traj models.

Since we found ballast SF-HON to be the best fitting model for the United States NIS risk and biofouling SF-HON is the most accurate model for Europe NIS risk, we provided a map of high-risk introduction pathways in Europe based on biofouling SF-HON and main introduction pathways in North America based on ballast SF-HON in figure S5.

### 3.5 Analysis of Different Vessel types

We further investigate how different vessel types contribute to ballast and biofouling NIS introduction risk. We realized that Bulk and Auto carriers are responsible for the majority of NIS introduction through ballast water. After that, Gas, Oil, and Passenger carriers are contributing the most, respectively. However, looking at table S2 we can see that 33.05% of the vessels in the ballast risk data are container carriers. However, they do not result in a high NIS introduction risk as Bulk carriers which are 20.21% of the records. Auto (8.53%), Gas (2.39%), Oil (8.53%), Passenger (2.01%) and carriers records (having high NIS introduction risk) are all less than General (15.83%) and Container (33.05%) carriers as well. This indicated that even though there are more Container and General carriers in the ballast risk records, they do not result in a high NIS introduction risk. We realized that based on our model, the reason is directly related to vessel Gross Weight Tonnage (GWT) which is a good measure of vessel size. Container and General carriers have a significantly smaller GWT compared to Bulk, Auto and Passenger carriers (Figure 8 (a)).

**Figure 8:**
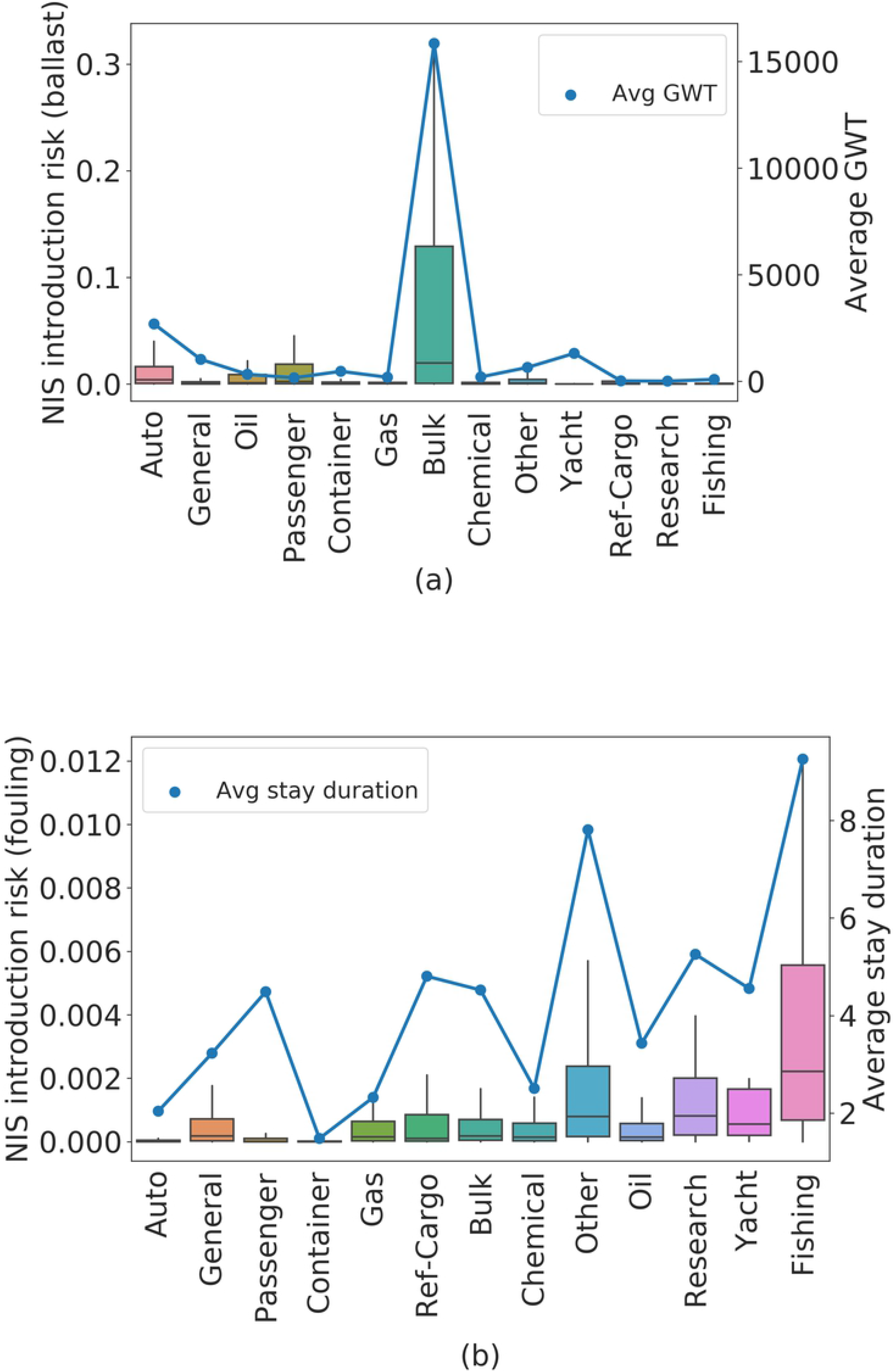
NIS introduction of different vessel types for ballast water (a) and biofouling (b). For ballast water, GWT and for biofouling, duration of stay at port display the same pattern as the NIS introduction risk.

We then performed the same analysis for the biofouling risk. We realized that in this case, many different carriers are responsible for introduction through biofouling. In particular, Fishing, Research, Other, and Yacht carriers are the main contributors to biofouling introduction. Again, table S2 indicates that these main contributes are not a large portion of all the risk records. Fishing carriers are only (0.17%) of the records. Research (0.13%), Yacht (0.08%), and Other (1.18%) containers each consist of a small portion of all the records while Container (23.07%), Bulk (21.68%), and General (21.68%) carriers are the major portion of the data. In this case, we realized that based on our model, the average duration of stay at source port displays the same pattern as the NIS introduction risk (figure 8 (b)). Recall that biofouling clusters are more localized. Many high-risk ports are distributed across these clusters. Fishing, Research, Yacht, (and possibly the “Other” category) correspond to several small vessels traveling through shorter distances which do not carry a large amount of cargo, and have fewer regulations imposed on them for using an antifouling system [28]. As a result, these local ships can easily transfer species locally within the small clusters in figure 4 (b). Therefore, vessel-based policies for ballast water prevention policies need to apply more stringent regulations for large vessels (Bulk and Auto carriers), while biofouling prevention policies need to target more vessel types and enforce more regulations for using an antifouling system for vessels that stay in harbors for a long time (Fishing, Research, Yacht, and “Other”)

## 4 Discussion

Analyzing both SF-HON models over 15 years highlights how the dynamics of shipping network can affect the NIS spread risk through the two main vessel introduction vectors. In particular, we realized that based on the proposed model, biofouling risk is highly affected by the duration of stay at the source port, while ballast transfer risk is mostly affected by the shipping traffic of the destination port. Our results indicate how a major change in world trade (e.g. the global recession) affected the shipping network and subsequent NIS spread risk. Our analysis provides insights for policymakers in terms of understanding the high-risk voyages, clusters of ports within which NIS spread risk is high, and targeting the inter-cluster connections or major hubs receiving diverse shipping traffic.

We revealed the importance of higher-order dependencies in modeling ship-borne NIS spread risk. We integrate global shipping, port environment, and bio-geographical datasets to build global risk models that include significant higher-order dependencies to construct SF-HONs based on both ship movements patterns and environmental conditions of the ports. Furthermore, our analysis for the first time models the biofouling risk using a world-wide species flow network model.

Our comparison of the SF-HONs and SF-FONs for both vessel vectors shows that higher-order patterns of species transfer are significantly more likely to occur through ballast water relative to biofouling. Furthermore, we show that SF-HONs predictions fit observed NIS introduction data better, than SF-FONs and existing higher-order model for species spread. This indicates that only significant higher-order dependencies need to be considered when modeling global ship-borne NIS spread to inform policy and management interventions aimed at reducing harmful NIS.

Our clustering analysis of ballast SF-HON and biofouling SF-HON reveals fundamental differences in species spread via each vessel vector, which indicates the necessity of imposing vector-based policies. Specifically, we emphasize on isolated biofouling clusters in which targeting the inter-cluster connection would be effective, versus several highly overlapping clusters in ballast SF-HON with significantly more higher-order ports and strong inter-cluster connections, which require local and global regulations and possibly targeting the high-risk ports with high shipping traffic.

Our analysis provides insights for policymakers in terms of understanding the high-risk voyages, clusters of ports within which NIS spread risk is high, and targeting the inter-cluster connections or major hubs receiving diverse shipping traffic. Furthermore, our work highlights the importance of vessel-based policies. In particular, such policies should vary with respect to the introduction vector. Ballast water prevention policies need to pose more restrictions on larger vessels, while biofouling prevention policies should target the vessels that stay in harbors longer.

Our treatment of the occurrence data is conservative. The NEMESIS and USGS data contain multiple records for many species. Because many of the records are probably results from secondary spread after an initial introduction by shipping, we included in our analysis only the chronologically first record. To the extent that some multiple records excluded from our analysis represent independent ship-related introductions, our dataset is incomplete and therefore lacking power. Nevertheless, the increased accuracy of SF-HONs over SF-FONs is apparent.

It is important to note that our models for ballast and biofouling introduction based on the availability of data at a global scale, and so they can be further improved if more accurate and detailed data about vessels and voyage-related introduction risks becomes available. Also, we did not consider ballast-water exchange and other ballast water management practices that were being practiced in some jurisdictions during some of our time series; such management actions could be simulated in future model versions. Finally, our analysis can be further expanded by breaking down the risk with respect to different temperature and salinity tolerance groups to identify the NIS spread pathways corresponding to specific species (e.g., [14]).

## Acknowledgements

This paper is based on research supported by the U.S. National Science Foundation Award number 1427157 (PIs: DM Lodge, NV Chawla, EK Grey), and the Army Research Laboratory under Cooperative Agreement Number W911NF-09-2-0053 (PI: NV Chawla). M.S., J.X., E.G., D.L., and N.C. collectively conceived the research. MS, JX, and E.G., designed the analyses. MS conducted the experiments. M.S., J.X., E.G., D.L. and N.C. wrote the manuscript. M.S. and J.X. contributed equally to the manuscript. We thank Salvatore Curasi for his valuable comments.

---

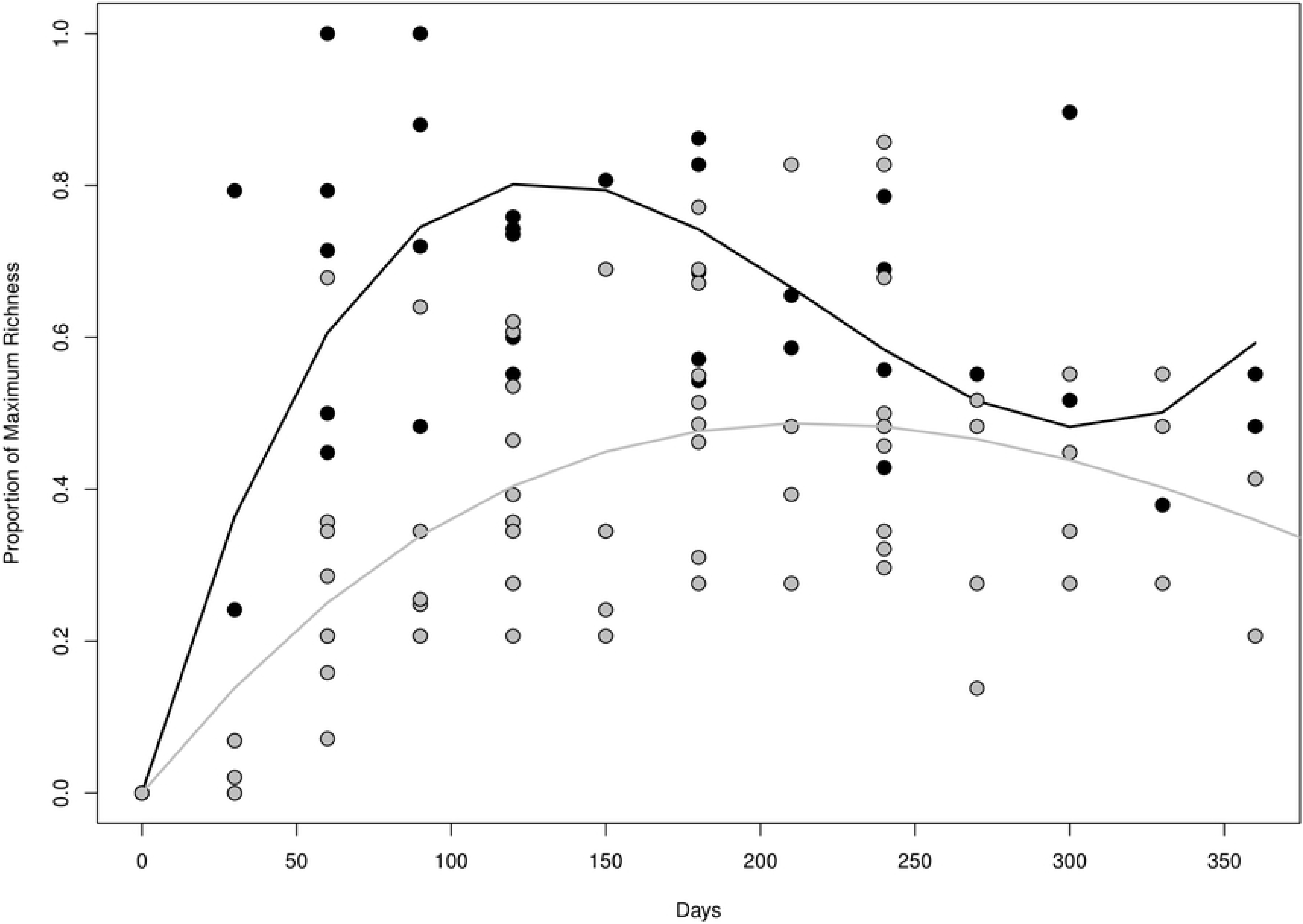

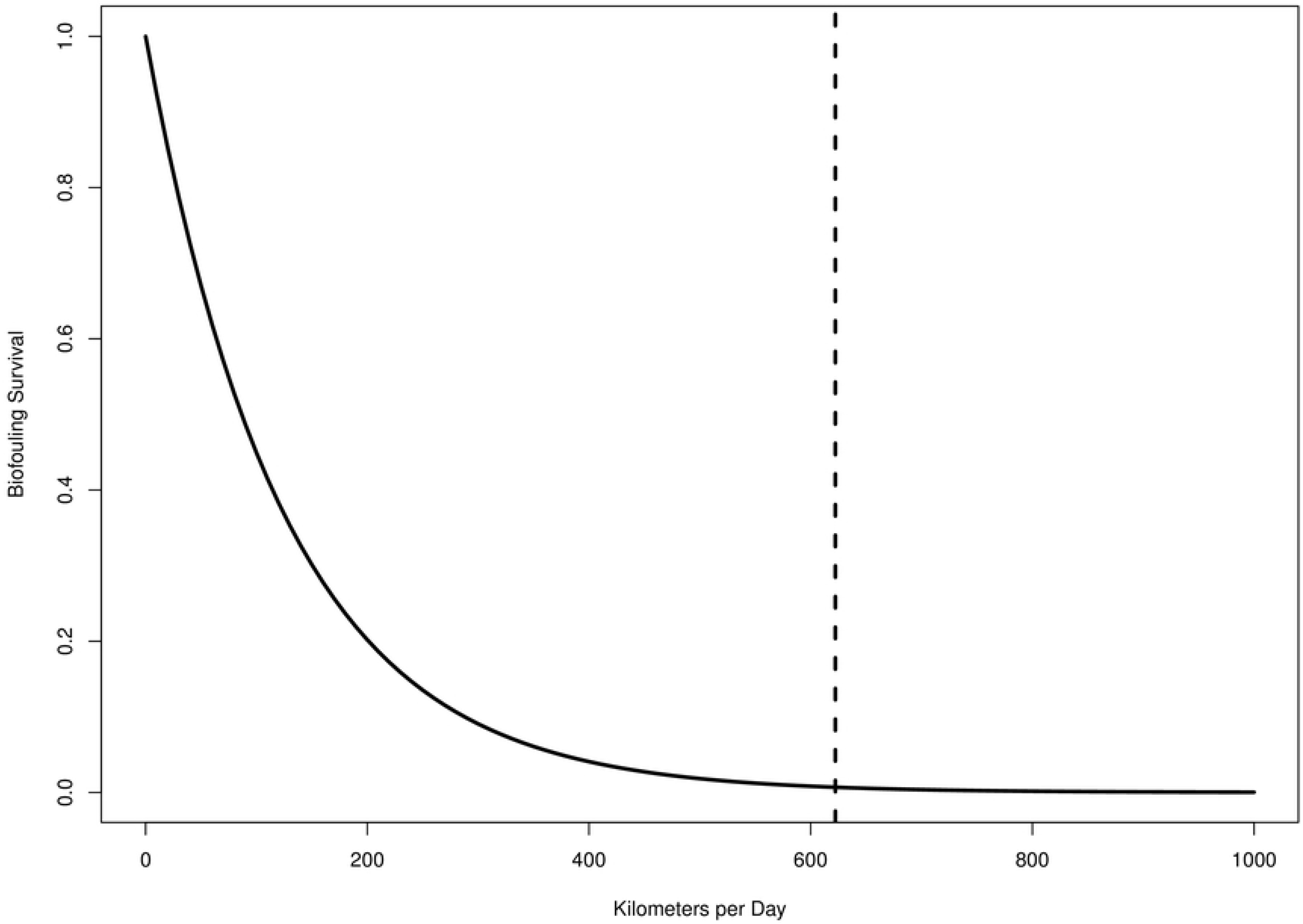

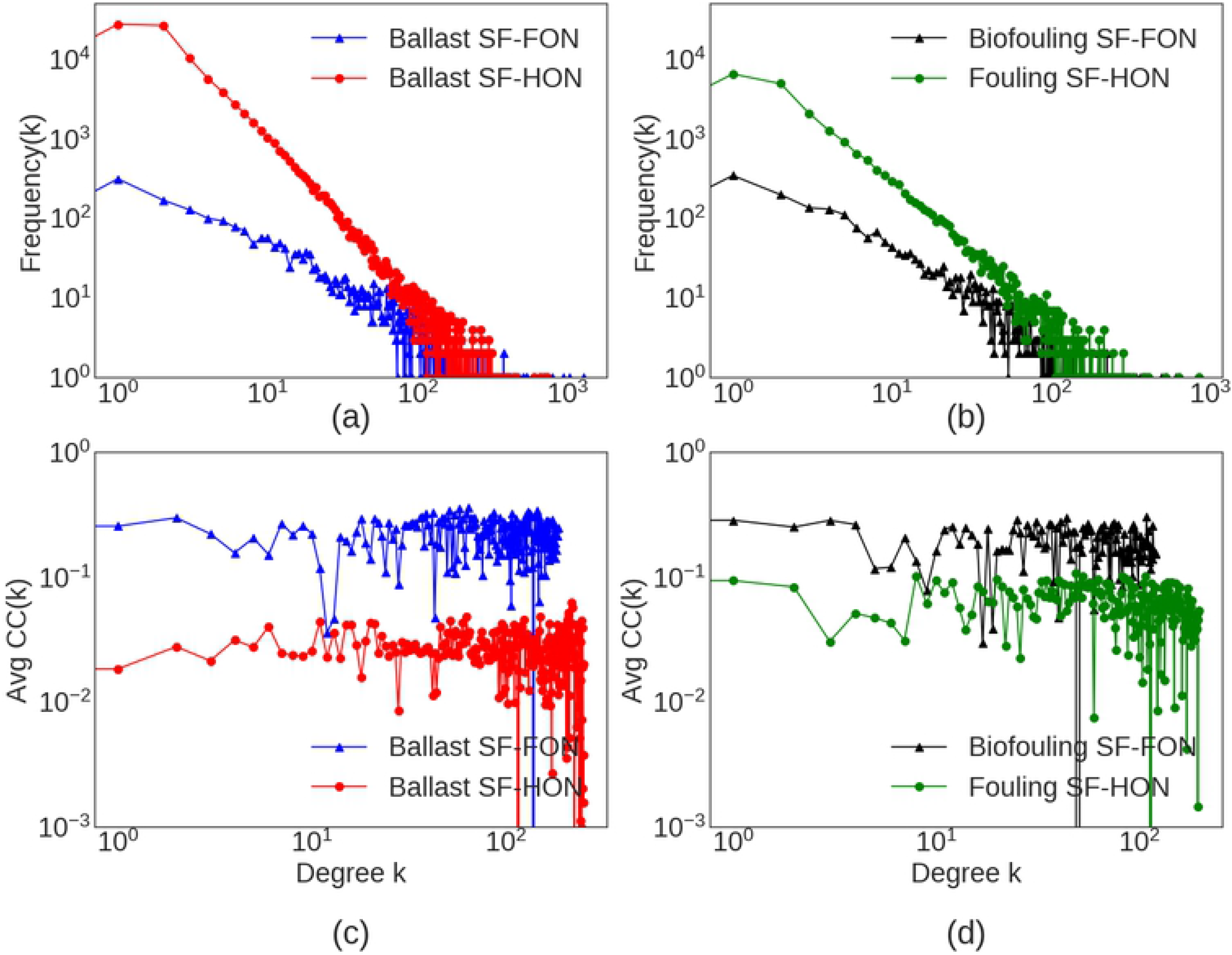

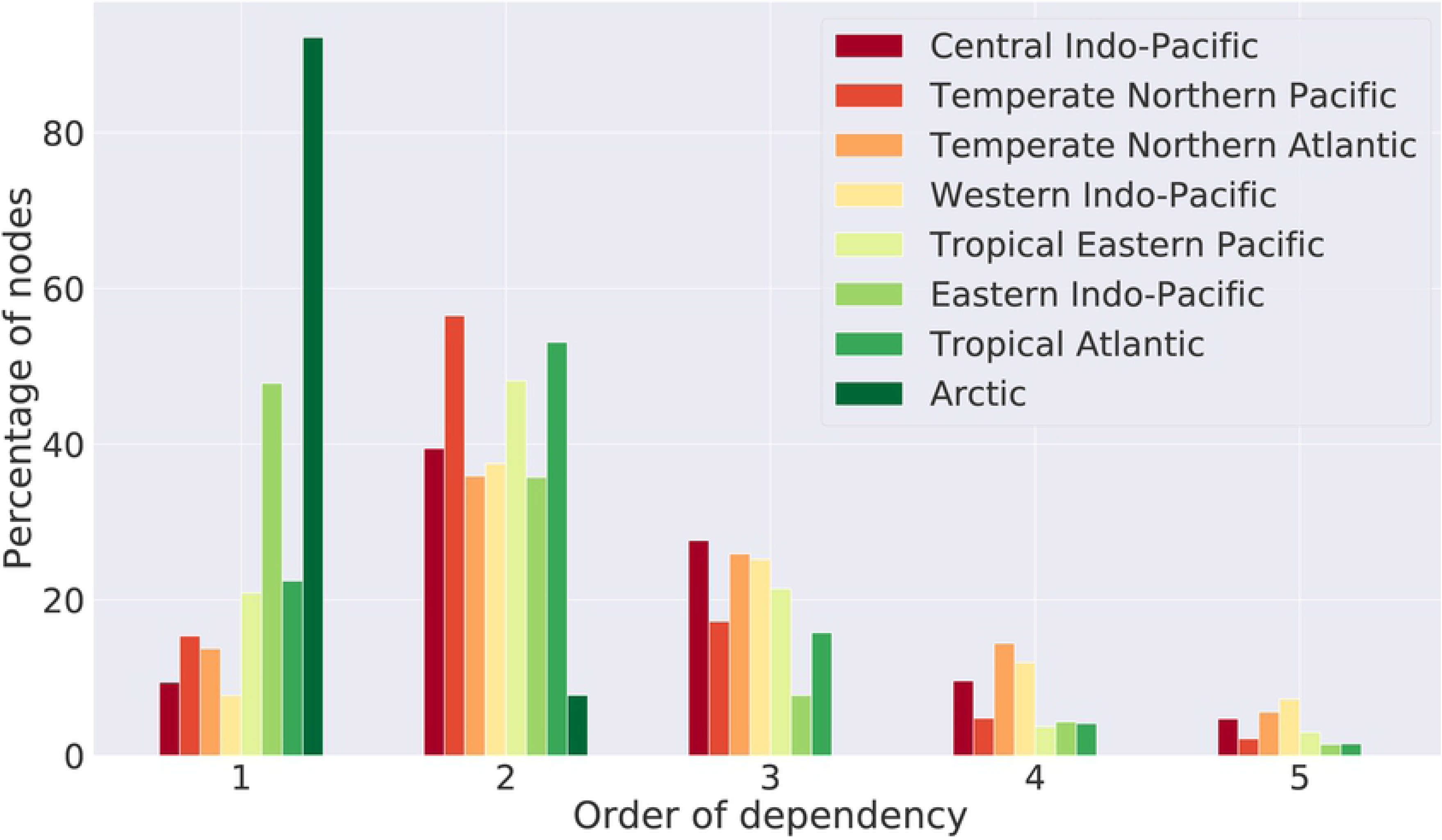

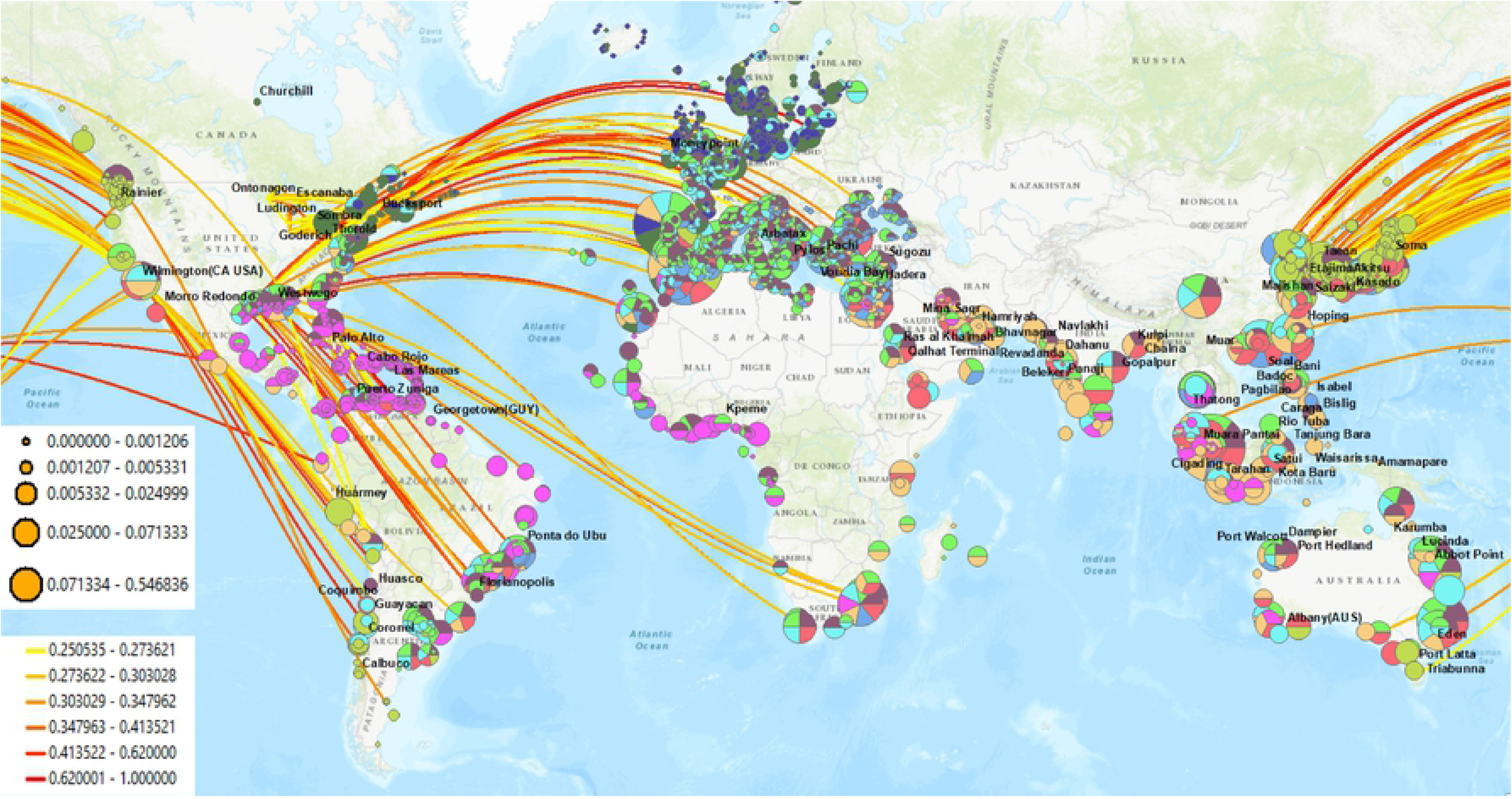
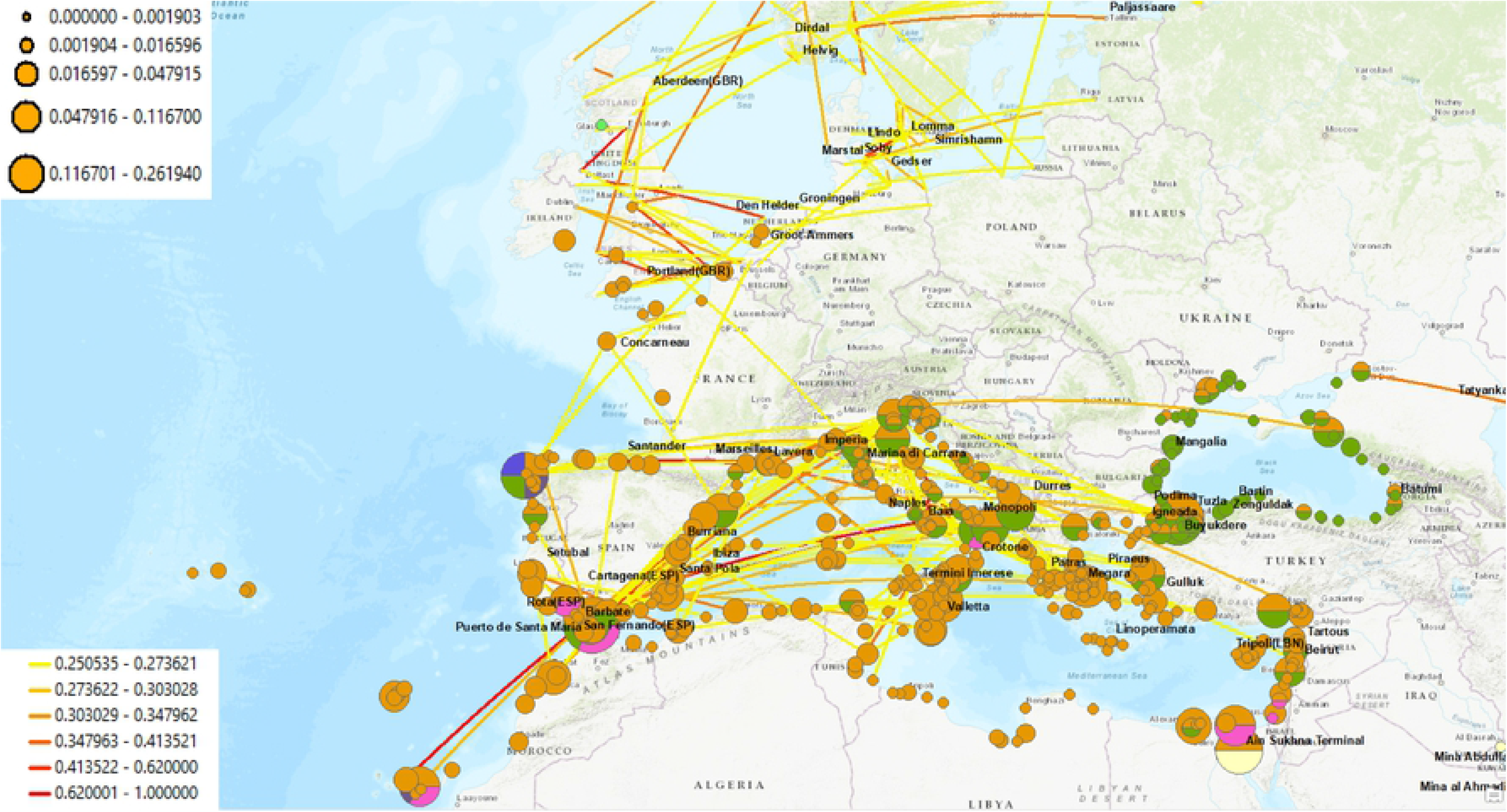

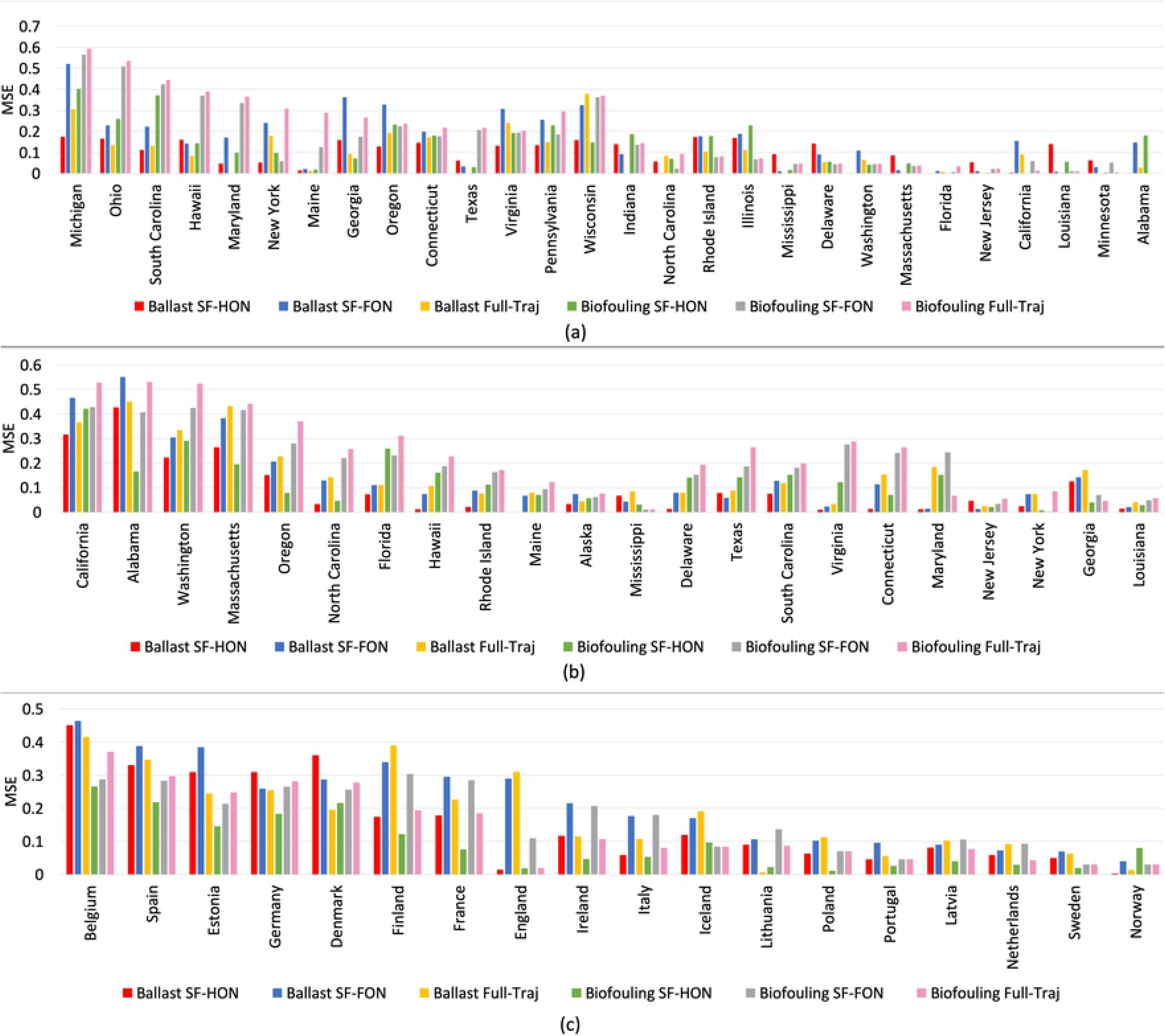

